# Graph-based Drug Decomposition for Anticancer Response Modeling

**DOI:** 10.64898/2026.01.06.697864

**Authors:** Stefano Nasini, Luis Fernando Pérez Armas, Oussama Bouaggad, Sophie Dabo, Djohana Laurent, Arnaud Petit, Meyling Cheok

## Abstract

This paper studies the molecular effects on *ex vivo* drug response in pediatric acute myeloid leukemia (AML). We firstly estimate dose–response relationships through linear and mixed-effects models, capturing both patient-specific heterogeneity and drug-level effects. Then, drug identifiers are decomposed into curated molecular descriptors and their higher-order interactions, yielding a structured, interpretable representation of chemical properties. To handle the resulting high-dimensional system, we introduce a specialized graph-based drug decomposition and selection, enabling a computationally tractable estimation of the molecular effects on *ex vivo* drug response. This strategy uncovers the molecular features most strongly associated with cellular viability, providing a biologically grounded and transparent alternative to black-box predictive methods. By directly linking molecular structure to therapeutic outcomes, our framework supports a novel mathematical-programming-based drug-combination selection and cost-efficient compound prioritization in pre-clinical leukemia research.

## 1 Introduction

AML is an aggressive and heterogeneous hematologic malignancy, the treatment of which is essentially based on chemotherapy. Despite substantial progress in therapeutic strategies, innovation in pediatric oncology has remained limited over the past two decades. One central reason is that research and development in this field often yields modest returns on investment, largely due to the low economic value associated with pediatric cancers, which account for only about 15% of cases in developed countries [Li et al., 2007]. A major obstacle lies in the design and execution of pre-clinical trials, an essential but costly component of the drug development process. These trials involve manufacturing and testing tens of thousands of potential compounds, yet fewer than 1% advance to clinical evaluation. This phase alone typically spans one to six years and accounts for costs ranging from $161 million to $4.54 billion [Schlander et al., 2021]. The resulting combinatorial explosion of candidate compounds severely undermines the financial viability of developing new therapies, particularly in pediatric oncology.

To mitigate this bottleneck, we study the molecular determinants of *ex vivo* drug response in pediatric AML through a graph-based decomposition and selection of existing anticancer drugs, with the aim of prioritizing promising compounds without replicating a large array of pre-clinical trials. The underlying idea is that, due to the complex molecular structure of anticancer drugs (which can be represented as graphs), their overall impact on pediatric AML can be decomposed into chemical substructures and physicochemical descriptors. To validate this approach, we leverage patients included in the MYECHILD01 and ALARM3 clinical trials (NCT02724163, NCT05772559), evaluated within the French CONECT-AML network^1^. Our cohort consists of 323 pediatric AML patients, with samples collected at diagnosis or relapse. Each sample was exposed in duplicate *ex vivo* to a median of 18 antileukemic drugs (range 2–23), with eight point serial dose-response observations per patient–drug pair. Viability was quantified via the MTT assay [Van Meerloo et al., 2011], which measures the enzymatic reduction of tetrazolium salts to formazan as a proxy for cellular viability, yielding a robust readout of drug-induced cytotoxicity. Rather than treating each compound as an indivisible label, our approach represents drugs through curated molecular descriptors and their higher-order interactions, organized as an interpretable graph. This structured representation is designed to connect observed response patterns to molecular constituents, thereby supporting mechanistic insight and principled compound selection in settings where exhaustive experimental screening is impractical.

From a methodological perspective, preclinical drug response analysis has increasingly relied on hierarchical statistical models, machine learning methods, and molecular graph-based representations. Hierarchical models offer an attractive balance between predictive performance and interpretability, enabling integration of pharmacological, genomic, and clinical covariates while distinguishing patient-level random effects from population-level fixed effects. For example, Cremaschi et al. [2024] propose a Bayesian hierarchical autoregressive model for residual disease dynamics in ALL, while Zeng et al. [2022] and Bottomly et al. [2022a] develop mixed models for AML drug response at single-cell or bulk levels. Similarly, White et al. [2021] and Li et al. [2024] introduce Bayesian hierarchical frameworks to disentangle drug sensitivity components and quantify tumorspecific molecular targets. Other contributions include active learning designs for compound prioritization [Tosh et al., 2025] and causal single-cell models of therapy response [Huang et al., 2024]. These works demonstrate the flexibility of hierarchical structures, but their scalability and clinical adoption remain constrained by computational cost, data heterogeneity, and interpretability.

In parallel, molecular graph representations have become a cornerstone of computational pharmacology. Graph-based signatures and chemical fragment decompositions capture essential molecular substructures, improving generalization across compounds [Cesar de Souza Costa et al., 2020, Rodrigues et al., 2021]. Applications span oncology [Evans et al., 2023, Janizek et al., 2021], toxicity prediction [Jiang et al., 2021b], and natural product discovery [Mullowney et al., 2023], with recent studies emphasizing explainable modeling frameworks [Lamens and Bajorath, 2023].

A closely related line of work benchmarks learned molecular representations, notably graph neural networks (GNNs), against classical descriptor-based models on standardized public benchmarks for molecular property prediction: Jiang et al. [2021a] compare multiple descriptor-based and graph-based architectures across a suite of widely used datasets (often referred to collectively as *MoleculeNet*), spanning both regression endpoints (e.g., physicochemical properties) and classification endpoints (e.g., bioactivity and toxicity labels). Their results highlight that descriptor-based models can remain highly competitive in predictive accuracy while being lighter and more directly interpretable, whereas GNN gains tend to emerge in specific settings and often at the cost of additional computational and modeling complexity. Importantly, the focus of such benchmarking studies is primarily predictive performance on molecule–endpoint datasets, rather than modeling patient- and dose-resolved *ex vivo* response or enabling downstream decision-making for compound selection in a clinical translational context. Yet, despite their predictive success, many graph-based approaches operate as opaque black-boxes, raising concerns in high-stakes clinical applications. As argued by Rudin [2019], intrinsically interpretable models should be prioritized over post-hoc explanations to ensure reliability and transparency in decision-making.

Building on these developments, our graph-informed statistical modeling framework for anticancer drug response in pediatric AML relies upon two complementary regression strategies: a drug effect model (DEM) and a molecular effect model (MEM). The DEM treats each compound as a fixed categorical effect, enabling individualized estimation across drugs but limiting the ability to extrapolate to novel compounds. By contrast, the MEM decomposes drug effects into molecular subgroup features, allowing us to link therapeutic efficacy to interpretable molecular constituents. In contrast to purely representation-learning approaches (e.g., end-to-end GNN predictors as surveyed and benchmarked by Jiang et al., 2021a), our MEM is intrinsically interpretable: drug effects are expressed directly in terms of curated chemical substructures and physicochemical descriptors and their higher-order interactions, while patient heterogeneity and concentration–response dynamics are modeled explicitly through the hierarchical regression structure. To handle the high dimensionality of the MEM decomposition, we introduce a Jacobi-based Lasso scheme with annealed penalization, enabling stable estimation while mitigating shrinkage bias, and yielding a computationally tractable estimation which bridges hierarchical regression with graph-based chemical representations and produces biologically grounded models of therapy response. In summary, our contribution lies in unifying hierarchical regression with graph-informed molecular decomposition for drug response modeling in pediatric AML. By explicitly linking therapeutic efficacy to interpretable structural features, the proposed DEM and MEM frameworks provide a transparent, extensible alternative to black-box prediction, supporting both compound prioritization in preclinical research and mechanistic insight into drug action.

Following this general introduction, Section 2 establishes the notation and definitions used throughout the paper and describes the DEM and MEM approaches. Section 3 focuses on our experimental design and data architecture. Sections 4 and 5 present the results of the DEM and MEM model, respectively.

## 2 Model

We consider a collection of *n* = 323 patients whose cancer cells are extracted and tested *ex vivo*. These measurements are used to test cancer cell resistance to a set 𝒟 of different drugs, with |𝒟| = *D* = 23, in our experimental design. In particular, 16 measurements (corresponding to 8 point serial drug dilution in duplicates) are made per patient–drug pair.^2^ In the rest of this paper, *j* indexes the patient and, for each fixed patient *j*, the index *i* enumerates all measured *ex vivo* well/plate conditions available for that patient (i.e., all drug–dose–replicate combinations recorded for *j*). In particular, *i* does not represent calendar time; it is simply an enumeration of experimental conditions within patient *j*. We denote by *d*(*i, j*) the drug index used for cell exposure under condition *i* for patient *j*, and by *W_i,j_* the corresponding cell viability, computed according to the standard formula:

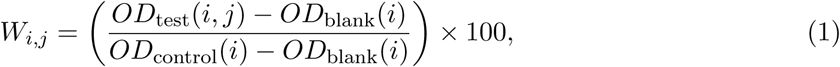

where *OD*_test_(*i, j*) is the optical density measured from the treated sample for patient *j* under condition *i*; *OD*_control_(*i*) is the mean OD from untreated control cells (representing 100% viability) under the same condition; and *OD*_blank_(*i*) is the mean OD from blank wells used to correct for background absorbance. Hence, *OD*_control_(*i*) and *OD*_blank_(*i*) are condition-specific calibration values shared across patients measured under the same plate condition *i*. The calculation of optical density is common in absorbance-based viability assays such as MTT, MTS, XTT, or WST-1 [Van Meerloo et al., 2011].

Our modeling formulation for quantifying anticancer therapy response defines a transformed viability outcome *Y_i,j_* = *ω*(*W_i,j_*) (based on well-defined transformations, as discussed in Section 4) and proposes two alternative regression approaches for the expected *Y_i,j_*: DEM and MEM. The DEM treats each compound as a fixed categorical effect, enabling individualized estimation across drugs but limiting the model’s ability to generalize to novel treatments or exploit shared functional molecular characteristics. Conversely, the MEM specification consists in projecting these compounds into a collection 𝒫 of molecular subgroup features (with |𝒫| = *P* = 57), which decompose the drug effects into chemical substructures and physicochemical descriptors. This choice contrasts with end-to-end representation learning approaches based on GNNs, which learn latent molecular embeddings optimized for prediction across benchmark molecular endpoints [Jiang et al., 2021a]. In our setting, the primary objective is not to learn a black-box embedding, but rather to obtain an *explicit*, clinically usable decomposition of drug identity into interpretable molecular factors that can be combined with patient heterogeneity and dose information.

### Regression models

Formally, the DEM and MEM approaches are expressed as follows:

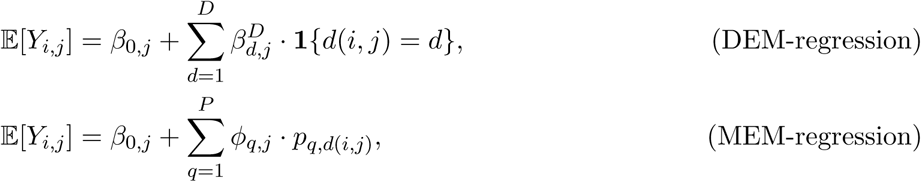

where *β*_0*,j*_ denotes the patient-specific intercept, and drug effects are interpreted relative to a chosen reference level (or under an equivalent identifiability constraint).

**Remark 1** (DEM terms). In (DEM-regression), the term 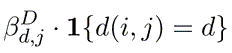 accounts for the identity of the administered drug, with 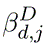 representing the effect of drug *d* on patient *j*, and **1**{*d*(*i, j*) = *d*} an indicator function selecting the relevant drug under condition i. Thus (DEM-regression) treats each compound as a fixed categorical effect.

**Remark 2** (MEM terms). In (MEM-regression), the term *ϕ_q,j_* · *p_q,d(i,j)_* captures the contribution of property *q*, when present in the administered drug *d*(*i, j*), to the expected viability outcome. In this notation *p_q,d_* ∈ {0, 1} indicates whether drug *d* exhibits molecular property *q*, and *ϕ_q,j_* represents the contribution of property q to the viability response in patient *j*. Under the MEM framework, the drug-specific effect 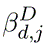 is expressed as a linear combination of its constituent properties:

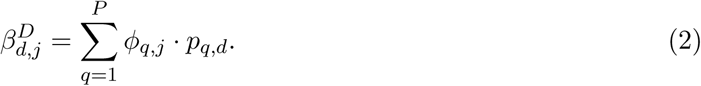

In this parameterization, the summation over discrete drug identifiers is replaced by a sum over molecular features. This enables out-of-sample prediction via linear extension over the property space, and supports mechanistic interpretation through the identification of salient molecular features associated with patient-specific treatment response.

Technically, (MEM-regression) can be viewed as a transparent alternative to the learned molecular representations compared in Jiang et al. [2021a]. In their setting, each molecule is mapped to a latent vector *z_d_* ∈ ℝ^*K*^ produced by a *GNN* (or to a fixed descriptor vector) and a downstream predictor is trained to map *z_d_* to an endpoint. By contrast, our representation *p*_·,*d*_ ∈ {0, 1}^*P*^ is fixed and human-interpretable, and the regression coefficients *ϕ_q,j_* (and their extensions to higher-order interactions in Section 5) directly quantify how specific substructures and physicochemical attributes contribute to patient-specific and dose-resolved response. This distinction is central for our downstream goal of principled compound and combination selection based on identifiable molecular mechanisms rather than on black-box embeddings.

### Biochemical covariates

In our empirical specifications, we allow both modeling formulations (DEM-regression) and (MEM-regression) to incorporate three biochemical covariates: the drug concentration 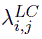, the optical density difference 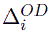, and the tumor/normal ratio 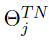. To account for biological and demographic heterogeneity, we further include patient-level covariates such as age and sex. The covariate 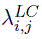 is included in (DEM-regression) and (MEM-regression) based on the following log-transformation: 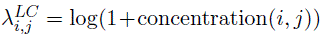, where concentration(*i, j*) refers to the concentration of drug under condition *i* for patient *j* (expressed in micromolar). The covariate 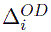 reflects the cells’ maximal metabolic capacity and is computed as 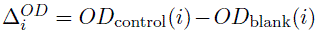, following the same measurement and reference-wavelength correction procedures described earlier. Here, *OD*_blank_(*i*) is a condition-specific calibration value (blank wells) defined at the well/plate condition level and therefore shared across patients measured under the same condition *i*; consequently, *OD*_blank_(*i*) does not carry a patient index. *OD*_control_(*i*) is specific to patient j under condition i and reflects no drug control. 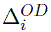. The covariate 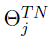 is based on the ratio of leukemic blasts to normal lymphocytes from the same sample and is an indicator of leukemia purity, potentially influencing cell viability. It is computed as the proportion of tumor cells divided by the proportion of normal cells.

## 3 Data

The dataset includes 323 pediatric AML samples recruited in France between 2018 and 2024, for which we performed an *ex vivo* drug sensitivity screen of primary blasts [Gonzales et al., 2022]. Briefly, 2 × 10^5^ blast cells per well were seeded in 96-well plates and exposed for 96 hours, under standard cell culture conditions, to an eight-point serial drug dilution in duplicates, resulting in 16 measurements per patient–drug pair. Our experimental design follows the MTT approach, where cellular metabolic activity is assessed by spectrophotometry (i3 Spectramax, Molecular Devices): mitochondrial enzymes reduce yellow MTT to violet formazan, and optical density (OD) is measured at 570 nm. The amount of formazan produced is proportional to the number of viable cells. For drugs showing absorbance at 570 nm, we correct *OD*_test_ by subtracting the corresponding drug-only control [de Haar-Holleman et al., 2004, Gonzales et al., 2022]. Table 1 reports summary statistics describing the resulting data architecture.

**Table 1:** Summary statistics for the principal experimental variables. Units are as follows: *W_i,j_* (cell viability) expressed as a percentage (%), 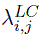 (log-transformed drug concentration) in (*µ*M), 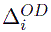 (optical density difference) unitless, and 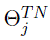 (tumour-to-normal ratio) unitless. Design parameters are reported in the lower portion of the table.

|  | Mean | SD |
| --- | --- | --- |
| $W_{i,j}$ | 53.715 | 47.088 |
| $\lambda_{i,j}^{LC}$ | 1.331 | 1.669 |
| $\Delta_i^{OD}$ | 0.263 | 0.223 |
| $\Theta_j^{TN}$ | 19.441 | 24.122 |
| Patients | $n = 323$ | |
| Samples per [drug, patient]-pair | $m = 8 \times 2$ | |
| Drugs | $D = 23$ | |
| Molecular properties | $P = 57$ | |

In principle, the cell viability score *W_i,j_* defined in (1) is bounded between 0% and 100%. However, we occasionally observe realized values of *W_i,j_*that are negative or above 100%. This can occur when *OD*_test_(*i, j*) *< OD*_blank_, for several reasons: (*i*) random measurement variability (typically on the order of ±10%), especially when the true signal (*OD*_control_) is low; (*ii*) strong cytotoxic effects that suppress metabolic activity to near, or below, the background level (i.e., *OD*_blank_ and the drug-only control); and (*iii*) overestimation of *OD*_blank_ due to technical artifacts. Negative values do not imply biologically “negative” viability; rather, they are consistent with complete or near-complete loss of viable cells relative to control, combined with experimental noise, and they are more frequent for colored compounds. Drug-only and cells-only controls are therefore important quality control measures. In statistical analyses, these values are either retained with their sign (to preserve the noise structure) or truncated at zero, depending on the modelling strategy. The list of *D* = 23 drugs is reported in Table 2.

**Table 2:**
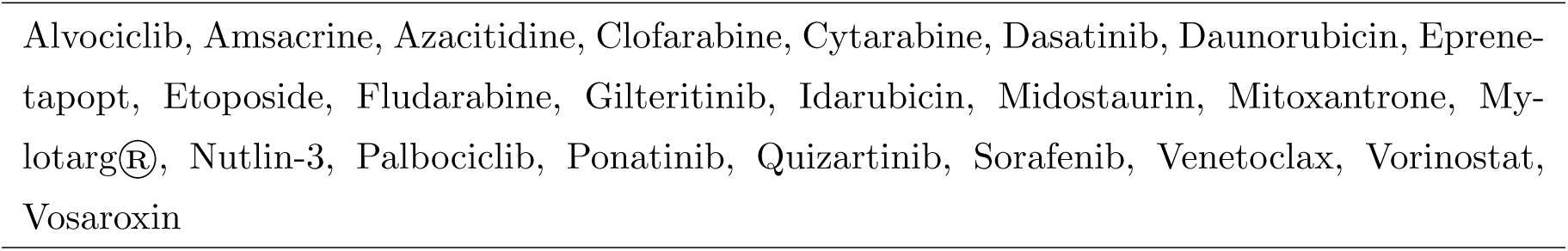
List of *D* = 23 drugs.

Figure 1 illustrates drug sensitivity profiles across the patient cohort using two complementary summary metrics.

**Figure 1:**
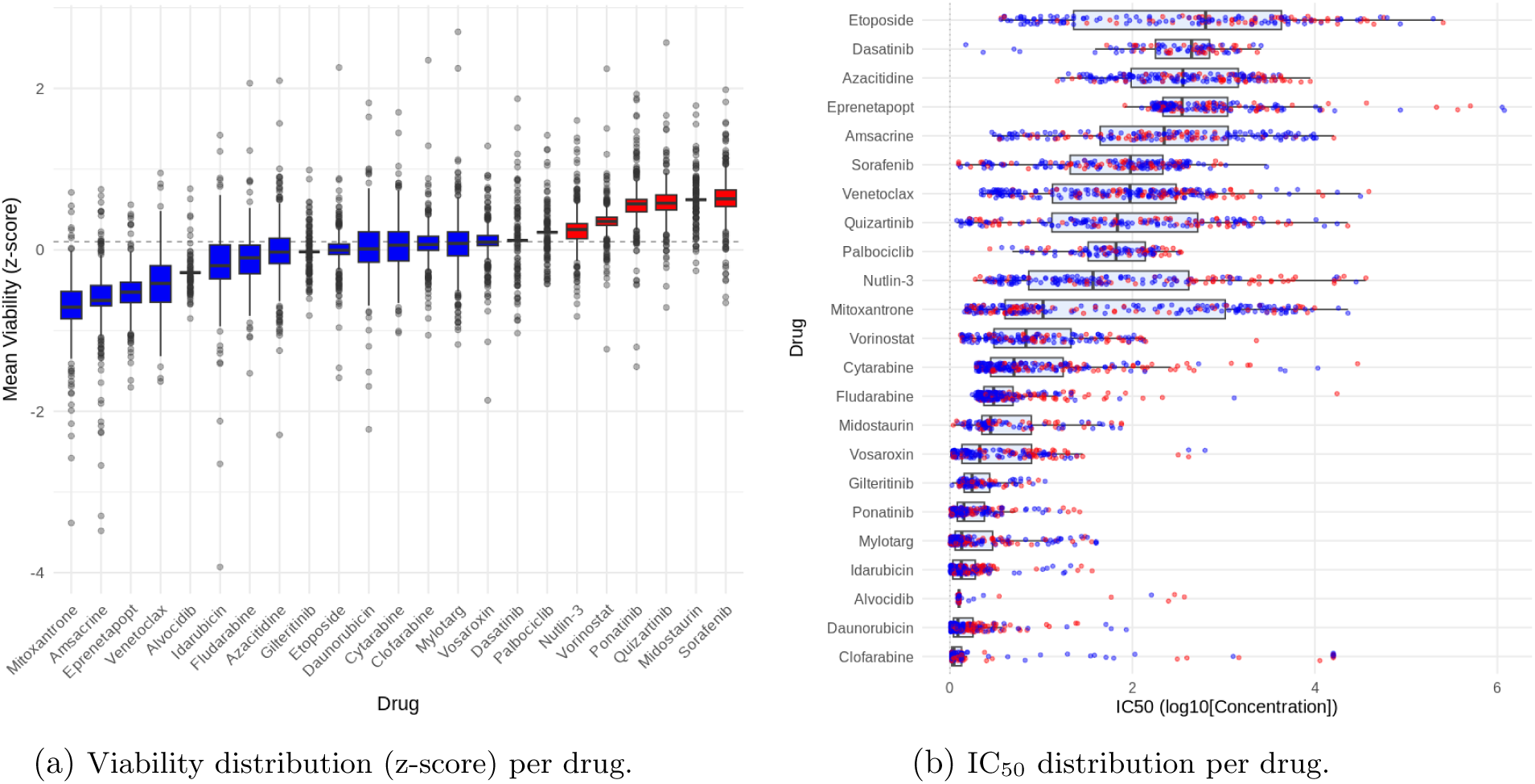
Drug response metrics across patients: (a) cell viability distribution (z-score) and (b) IC_50_ distribution (log_10_ scale). Drugs are ordered by their median response, with consistent patterns distinguishing sensitive (blue) versus resistant (red) compounds.

Panel (a) of Figure 1 summarizes *ex vivo* sensitivity by displaying, for each drug, the distribution of standardized viability scores (z-scores) across patients. Drugs are ordered by their median z-score and labeled as sensitive (blue) or resistant (red) according to whether this median lies below or above the prespecified threshold (dashed line); more negative medians indicate stronger overall cytotoxic effects (lower viability). Panel (b) reports the corresponding distribution of half-maximal inhibitory concentrations (IC∗50) across patients (log∗10 scale, in *µ*M). Smaller median IC_50_ values reflect higher potency, as lower concentrations are sufficient to reduce viability by 50%.

Focusing on the *D* = 23 drugs, we consider *P* = 57 molecular properties, selected to capture recurrent topological substructures and pharmacological patterns, as reported in Table 3. These 57 properties are grouped according to whether they are (i) purely topological substructures that can be computed symbolically from molecular graphs (first group), or (ii) pharmacological and physicochemical attributes that typically rely on empirical characterization and database curation (second group). For the first group, we assess graph similarity using the maximal common substructure (MCS) approach, as implemented in the Python library RDKit [Scalfani et al., 2022]. Each compound is represented by its canonical SMILES string and converted into a molecular graph object, from which we compute the MCS between each drug and a library of reference scaffolds (e.g., purine, pyrimidine, naphthalene, indole, ribose, etc.). This allows us to identify whether a given substructure is present in the drug molecule and to encode conserved aromatic cores or heterocyclic frameworks as molecular descriptors. In addition, RDKit provides atom- and bond-level features such as element counts (#C, #N, #O, halogens), stereochemical descriptors (defined/undefined atom and bond stereocenters), and topological indices (ring count, rotatable bond count, hydrogen bond donors/acceptors, heavy atom count). These properties are computed directly from the graph representation and stored as binary or numeric indicators. For the second group, properties are obtained from standard chemical annotations reported in major pharmaceutical databases such as DrugBank and PubChem.

**Table 3:**
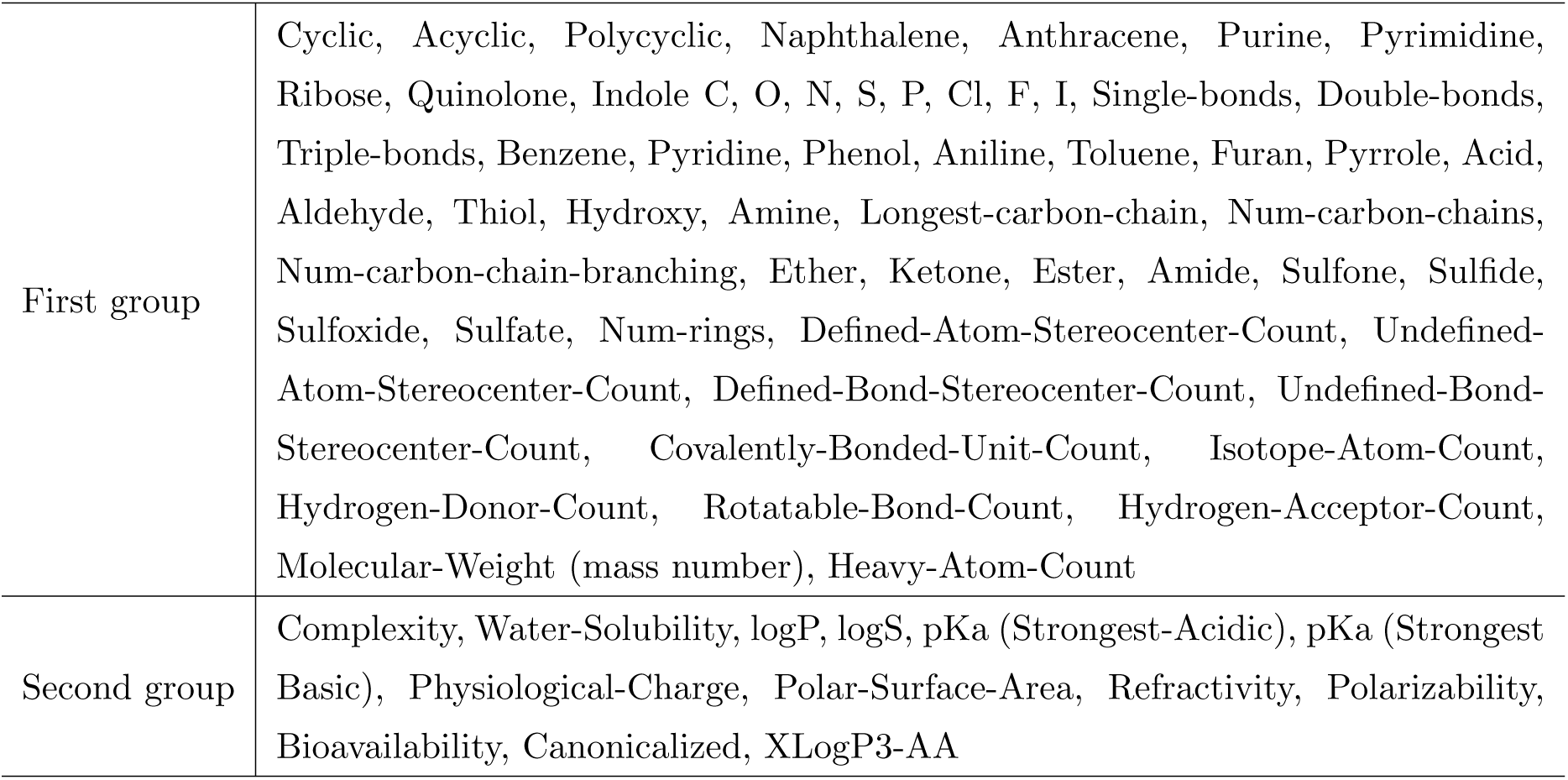
List of *P* = 57 molecular properties.

|  |  |
| --- | --- |
| First group | Cyclic, Acyclic, Polycyclic, Naphthalene, Anthracene, Purine, Pyrimidine, Ribose, Quinolone, Indole C, O, N, S, P, Cl, F, I, Single-bonds, Double-bonds, Triple-bonds, Benzene, Pyridine, Phenol, Aniline, Toluene, Furan, Pyrrole, Acid, Aldehyde, Thiol, Hydroxy, Amine, Longest-carbon-chain, Num-carbon-chains, Num-carbon-chain-branching, Ether, Ketone, Ester, Amide, Sulfone, Sulfide, Sulfoxide, Sulfate, Num-rings, Defined-Atom-Stereocenter-Count, Undefined-Atom-Stereocenter-Count, Defined-Bond-Stereocenter-Count, Undefined-Bond-Stereocenter-Count, Covalently-Bonded-Unit-Count, Isotope-Atom-Count, Hydrogen-Donor-Count, Rotatable-Bond-Count, Hydrogen-Acceptor-Count, Molecular-Weight (mass number), Heavy-Atom-Count |
| Second group | Complexity, Water-Solubility, logP, logS, pKa (Strongest-Acidic), pKa (Strongest Basic), Physiological-Charge, Polar-Surface-Area, Refractivity, Polarizability, Bioavailability, Canonicalized, XLogP3-AA |

To clarify the relationship between the collection of drugs and their molecular properties, Figure 2 illustrates two representative compounds: fludarabine (Panel (a)) and daunorubicin (Panel (b)). For fludarabine, we highlight its association with the purine scaffold, while for daunorubicin we report its link to the anthracene scaffold. The MCS between purine (bottom of Panel (a) of Figure 2) and fludarabine (top of Panel (a) of Figure 2) indicates that the purine scaffold is fully present in fludarabine, consistent with its ability to mimic natural nucleotides leading to its incorporation into nucleic acids, but then causing DNA double strand breaksand cell death. Similarly, the MCS between daunorubicin (top of Panel (b)) and anthracene (bottom of Panel (b)) indicates the presence of a planar anthracene tricyclic aromatic core, which is critical for DNA intercalation and the disruption of replication and transcription.

**Figure 2:**
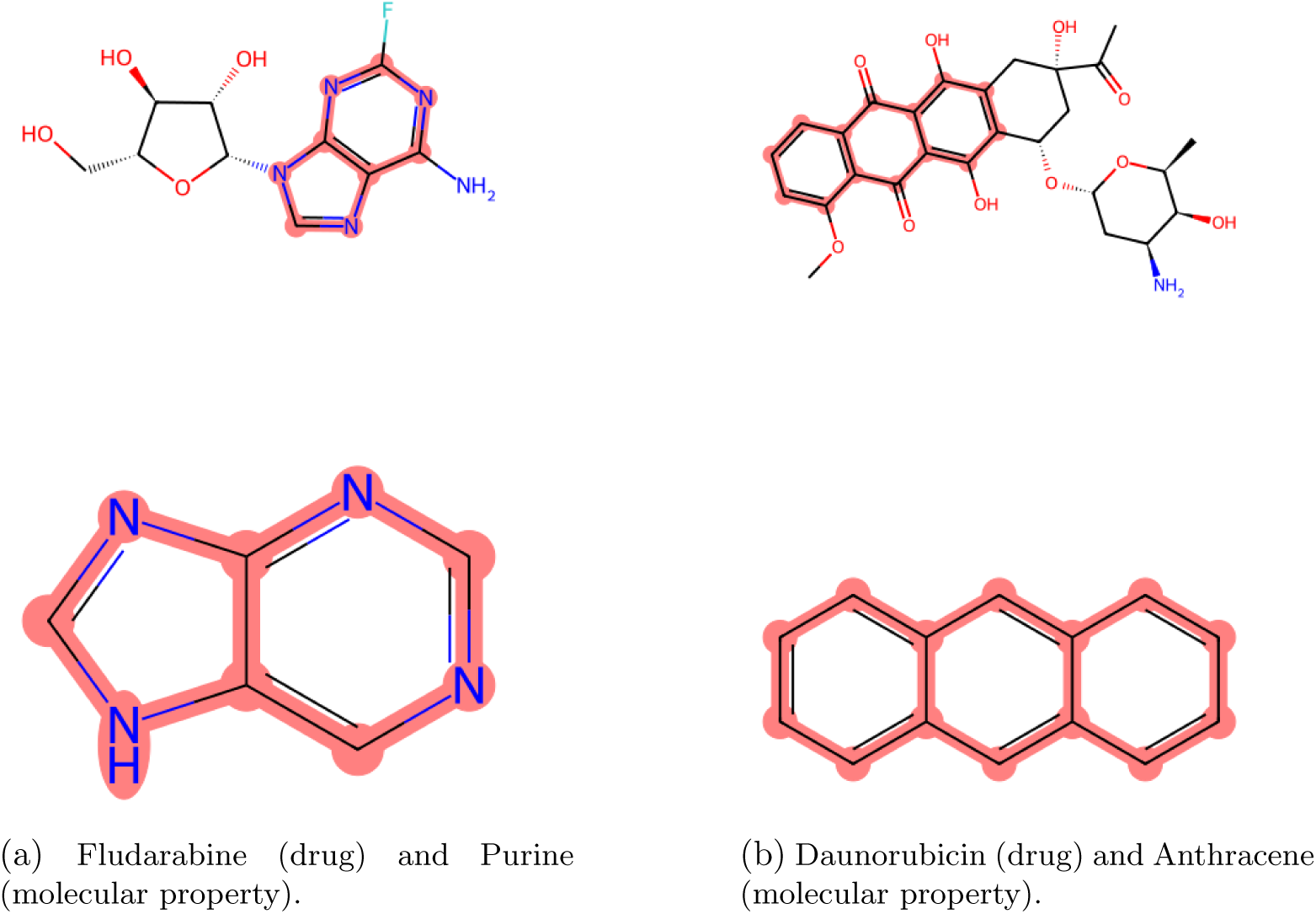
Structural comparison of drug molecules with their parent scaffolds. In Panel (a), Fludarabine (top) contains the purine heterocyclic framework (bottom), which supports its capacity to interfere with nucleotide metabolism and DNA synthesis. In Panel (b), Daunorubicin (top) shares the anthracene core (bottom), enabling DNA intercalation; appended functional groups modulate activity and pharmacokinetics.

An important constituent of our data architecture is drug concentration, which has a direct effect on the viability score. Figure 3 illustrates the aggregate dose–response trajectory across all drugs and patients. The smooth decreasing trend in standardized viability confirms the expected monotonic relationship between drug concentration and cytotoxicity: as doses increase, cell viability declines. The relatively narrow standard error bands highlight consistent responses across drugs despite baseline heterogeneity, supporting the experimental design and the subsequent regression analyses, where concentration is consistently identified as a dominant predictor of viability.

**Figure 3:**
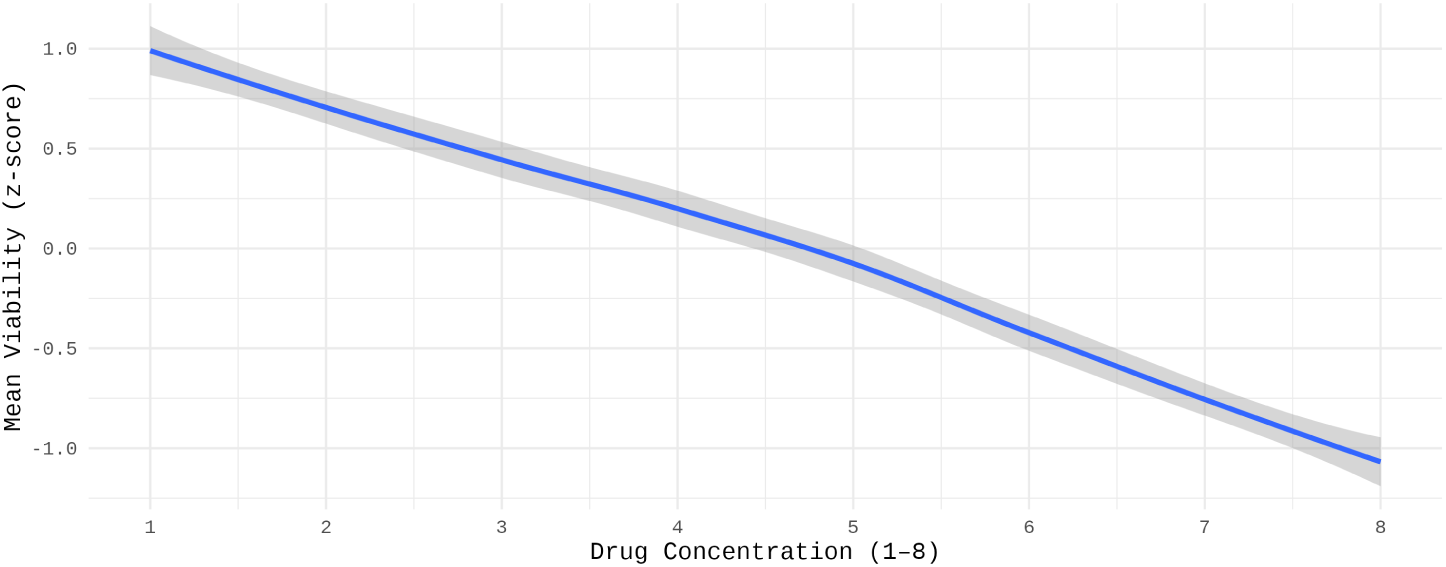
Generalized LOESS-smoothed trajectory of standardized viability (*z*-score) across the eight dose levels (obs ∈ {1, …, 8}). Each point represents the mean ± SE over all drugs and patients after per-drug normalization, highlighting the aggregate dose–response pattern independent of baseline viability scales.

## 4 Results on the DEM

In this section, we consider four specifications of the DEM regression in (DEM-regression), which differ by (i) the transformation *ω* applied to the viability response and (ii) the way patient effects are modeled. The first specification uses no transformation (*ω*(*x*) = *x*), assumes a Gaussian response for *W_i,j_*, and includes patient fixed effects *β*_0*,j*_ for each *j* ∈ {1*, …, n*}. The second specification also uses no transformation and a Gaussian response model, but treats the patient intercepts *β*_0*,j*_ as random effects drawn from a common Gaussian distribution. The third specification applies a min–max transformation *ω* and models *ω*(*W_i,j_*) as log-Gaussian; it also assumes Gaussian patient random effects for *β*_0*,j*_.

We first consider a standard linear model (LM) with patient fixed effects and estimate patient- and drug-specific concentration–response effects, using two transformations of the original viability measurements *W_i,j_* ∈ ℝ defined in (1). Specifically, we define the transformed outcome *V_i,j_*= *ω*(*W_i,j_*) as

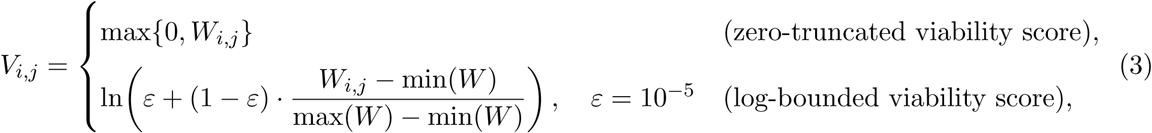

where min(*W*) and max(*W*) are taken over the observed viability values in the dataset. The regression model corresponds to an augmented version of (DEM-regression), in which we include the log-transformed concentration covariate 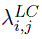 and interaction terms allowing patient-specific and drug-specific concentration effects, together with fixed effects for the administered drug **1**{*d*(*i, j*) = *d*} and for patients. Tables 4 and 5 report, respectively, the estimated drug fixed effects and a summary of the patient- and drug-specific concentration effects. The corresponding *R*^2^ coefficients (based on the *n* = 323 patient dataset described in Section 3) are reported at the bottom of Table 4.

**Table 4:** Fixed effect estimates and *p*-values for both the zero-truncated and bounded viability scores based on a patient-fixed-effect model. Both models are fitted without an intercept and include interaction terms encoding patient-specific and drug-specific log-concentration coefficients 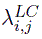.

| Term | Zero-truncated viability score |  | Log-bounded viability score |  |
| --- | --- | --- | --- | --- |
|  | Estimate | p-val | Estimate | p-val |
| (Intercept) | 42.12680 | < 2e-16 *** | -0.6264 | < 2e-16 *** |
| Amsacrine | 2.01887 | 0.000877 *** | -0.006252 | 8.56e-07 *** |
| Cytarabine | 18.72029 | < 2e-16 *** | 0.01607 | < 2e-16 *** |
| Azacitidine | 52.79273 | < 2e-16 *** | 0.04829 | < 2e-16 *** |
| Binimetinib | 13.97541 | 1.15e-11 *** | 0.01217 | 0.004751 ** |
| BMH21 | 2.48246 | 0.715134 | 0.000127 | 0.992861 |
| Clofarabine | -3.78964 | 9.93e-09 *** | -0.003483 | 0.011848 * |
| Dasatinib | 12.95667 | < 2e-16 *** | 0.01053 | 1.05e-11 *** |
| Daunorubicin | -15.90936 | < 2e-16 *** | -0.02472 | < 2e-16 *** |
| Etoposibe | 30.26883 | < 2e-16 *** | 0.02563 | < 2e-16 *** |
| Alvociclib | -16.30634 | < 2e-16 *** | -0.01508 | < 2e-16 *** |
| Fludarabine | 13.62876 | < 2e-16 *** | 0.01186 | < 2e-16 *** |
| Gilteritinib | -12.22909 | < 2e-16 *** | -0.01075 | 8.91e-11 *** |
| Homoharringtonin | -45.05222 | < 2e-16 *** | -0.04192 | 2.78e-12 *** |
| Idarubicin | -22.17295 | < 2e-16 *** | -0.02216 | < 2e-16 *** |
| MI3454 | 24.10402 | < 2e-16 *** | 0.02076 | 1.71e-07 *** |
| Midostaurin | 21.50449 | < 2e-16 *** | 0.01858 | < 2e-16 *** |
| Mitoxantrone | -13.83139 | < 2e-16 *** | -0.02299 | < 2e-16 *** |
| Mylotarg | 3.66560 | 1.41e-07 *** | 0.003858 | 0.008120 ** |
| Nutlin-3a | 30.57809 | < 2e-16 *** | 0.02623 | < 2e-16 *** |
| Palbociclib | 29.98339 | < 2e-16 *** | 0.02619 | < 2e-16 *** |
| Ponatinib | 4.16806 | 7.81e-08 *** | 0.003137 | 0.053404 . |
| Eprenetapopt | 49.59161 | < 2e-16 *** | 0.04364 | < 2e-16 *** |
| Quizartinib | 38.43087 | < 2e-16 *** | 0.03368 | < 2e-16 *** |
| Revumenib | 43.82199 | < 2e-16 *** | 0.03840 | < 2e-16 *** |
| Sorafenib | 33.95614 | < 2e-16 *** | 0.02930 | < 2e-16 *** |
| Trametinib | -6.21750 | 0.508587 | -0.006750 | 0.731700 |
| Voreloxin | -3.12128 | 4.81e-06 *** | -0.004038 | 0.004706 ** |
| Vorinostat | 18.08338 | < 2e-16 *** | 0.01563 | < 2e-16 *** |
| | $R^2 = 0.646$, Adj. $R^2 = 0.643$ | | $R^2 = 0.284$, Adj. $R^2 = 0.278$ | |

**Table 5:** Summary of patient- and drug-specific log-concentration effects 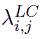a under the LM specifications.

| Term | Zero-truncated viability score |  | Log-bounded viability score |  |
| --- | --- | --- | --- | --- |
|  | Estimate | p-val | Estimate | p-val |
| mean (328) $\lambda_{i,j}^{LC}$ | -6.347 | 0.0055 ** | -0.0058 | 0.0199 ** |
| # p-values < 0.05 | 323 |  | 311 |  |
| # p-values < 0.01 | 319 |  | 296 |  |
| # p-values < 0.001 | 317 |  | 281 |  |

Tables 4 and 5 indicate a strong and consistent drug signal after accounting for patient heterogeneity and concentration effects. Across both response scalings (zero-truncated and log-bounded), most drug fixed effects are highly significant, with coefficients of varying sign: negative estimates correspond to reduced viability (stronger cytotoxic activity), while positive estimates correspond to higher viability (relative resistance) on the chosen response scale. Although absolute magnitudes differ because the two outcomes live on different scales, the overall significance patterns are broadly concordant. The in-sample fit is higher for the zero-truncated specification (*R*^2^ ≈ 0.646) than for the log-bounded specification (*R*^2^ ≈ 0.284). Turning to the concentration effects in Table 5, the mean patient- and drug-specific log-concentration coefficient is negative on both scales (zero-truncated: *λ^LC^* = −6.347, *p* = 0.0055; log-bounded: *λ^LC^* = −0.0058, *p* = 0.0199), indicating that, on average, higher drug concentrations reduce viability. Moreover, the vast majority of these *λ^LC^* coefficients are individually significant (zero-truncated: 323/328 at *p <* 0.05; log-bounded: 311/328 at *p <* 0.05), highlighting pronounced patient- and drug-specific concentration–response heterogeneity.

Overall, the results in Tables 4 and 5 (and the supporting multicollinearity check in Table A.1) suggest strong, drug-dependent reductions in viability with increasing concentration, while revealing substantial between-patient variability in the strength of this relationship.

To further account for this variability, we complement the LM analysis with a hierarchical linear model (HLM) in which the transformed viability score *V_i,j_* is modeled as Gaussian and patient heterogeneity is captured through random effects. The model is

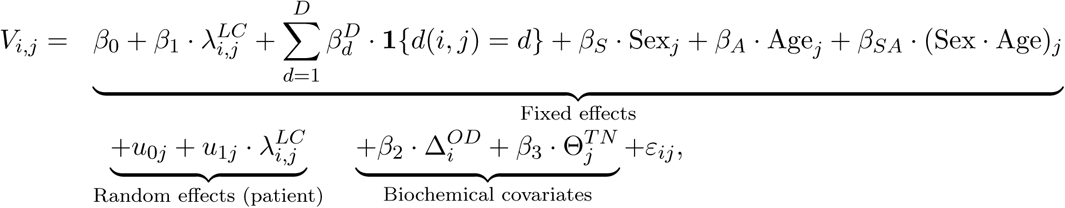

where *ε_ij_*~ 𝒩 (0*, σ*^2^) is an observation-level residual term, assumed independent across measurements. The patient-specific random intercept and slope, *u*_0*j*_ and *u*_1*j*_, are modeled jointly as a bivariate normal distribution:

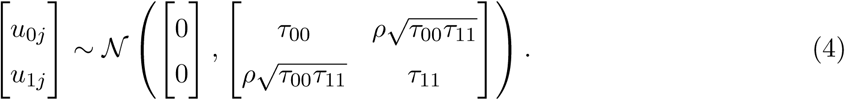

Table 6 reports the fitted fixed effects for this hierarchical model.

**Table 6:** Random effect estimates and *p*-values for both the zero-truncated and Log-bounded viability scores based on a patient-fixed-effect model. Both models are fitted without an intercept and include interaction terms encoding patient-specific and drug-specific log-concentration coefficients 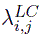.

| Term | Zero-truncated viability score |  | Log-bounded viability score |  |
| --- | --- | --- | --- | --- |
|  | Estimate | p-val | Estimate | p-val |
| (Intercept) | 42.61 | 0.000518 *** | -0.6318 | < 2e-16 *** |
| SexF | 3.921 | 0.746181 | -0.004418 | 0.743029 |
| SexM | 4.548 | 0.707078 | -0.003036 | 0.821574 |
| Age | 0.1646 | 0.163186 | 0.0001947 | 0.130420 |
| Sample.typeIF | -5.731 | 0.005365 ** | -0.001273 | 0.717461 |
| Sample.typeRech | -2.659 | 4.58e-08 *** | -0.003682 | 4.24e-05 *** |
| Ratio_TN | -0.05494 | 2.85e-05 *** | 0.0001249 | 2.46e-09 *** |
| Delta OD | 4.396 | 0.005533 ** | 0.01516 | 1.76e-09 *** |
| Amsacrine | 3.084 | 4.58e-06 *** | -0.006737 | 1.17e-07 *** |
| Cytarabine | 21.34 | < 2e-16 *** | 0.01570 | < 2e-16 *** |
| Azacitidine | 62.77 | < 2e-16 *** | 0.04837 | < 2e-16 *** |
| Binitinib | 12.91 | 8.08e-09 *** | 0.01161 | 0.007032 ** |
| BMH21 | 8.649 | 0.176078 | -0.0004659 | 0.973901 |
| Clofarabine | -3.948 | 3.19e-09 *** | -0.003740 | 0.006966 ** |
| Dasatinib | 11.24 | < 2e-16 *** | 0.009994 | 1.23e-10 *** |
| Daunorubicin | -15.23 | < 2e-16 *** | -0.02502 | < 2e-16 *** |
| Etoposide | 32.52 | < 2e-16 *** | 0.02541 | < 2e-16 *** |
| Alvociclib | -12.06 | < 2e-16 *** | -0.01539 | < 2e-16 *** |
| Fludarabine | 17.78 | < 2e-16 *** | 0.01167 | < 2e-16 *** |
| Gilteritinib | -3.932 | 3.20e-08 *** | -0.01079 | 8.34e-11 *** |
| Homoharringtonin | -40.96 | < 2e-16 *** | -0.04184 | 2.88e-12 *** |
| Idarubicin | -22.48 | < 2e-16 *** | -0.02229 | < 2e-16 *** |
| MI3454 | 28.14 | < 2e-16 *** | 0.02013 | 3.65e-07 *** |
| Midostaurin | 21.60 | < 2e-16 *** | 0.01825 | < 2e-16 *** |
| Mitoxantrone | -14.80 | < 2e-16 *** | -0.02337 | < 2e-16 *** |
| Mylotarg | 1.496 | 0.026838 * | 0.003698 | 0.011274 * |
| Nutlin-3a | 33.61 | < 2e-16 *** | 0.02581 | < 2e-16 *** |
| Palbociclib | 37.25 | < 2e-16 *** | 0.02626 | < 2e-16 *** |
| Ponatinib | 4.464 | 3.01e-09 *** | 0.002744 | 0.091762 . |
| Eprenetapopt | 56.83 | < 2e-16 *** | 0.04300 | < 2e-16 *** |
| Quizartinib | 36.58 | < 2e-16 *** | 0.03311 | < 2e-16 *** |
| Revumetinib | 49.64 | < 2e-16 *** | 0.03822 | < 2e-16 *** |
| Sorafenib | 34.20 | < 2e-16 *** | 0.02894 | < 2e-16 *** |
| Trametinib | -10.69 | 0.207673 | -0.007344 | 0.709339 |
| Voreloxin | -1.401 | 0.038777 * | -0.004476 | 0.001770 ** |
| Vorinostat | 21.01 | < 2e-16 *** | 0.01520 | < 2e-16 *** |
| | Condi. $R^2 = 0.650$, Marg. $R^2 = 0.567$ | | Condi. $R^2 = 0.293$, Marg. $R^2 = 0.249$ | |

The effect of log concentration (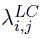) is strongly negative (Estimate = −0.396, *p <* 2 × 10^−16^), confirming that increasing concentration reduces cell viability (Figure 3). The tumor-to-normal ratio (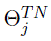) has a significant negative effect (Estimate = −0.05494, *p* = 2.85e−05), indicating that higher leukemia purity is associated with lower measured viability (i.e., stronger apparent drug effect). In contrast, the optical density difference (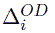) has a significant positive effect (Estimate = 4.396, *p* = 0.00553), suggesting that samples with higher metabolic capacity exhibit slightly higher viability. Neither sex nor age is significant, although their interaction is marginal (*p* ≈ 0.06). Most drug coefficients are highly significant, with negative estimates corresponding to lower viability (stronger cytotoxicity) on the modeled scale. In summary, the numerical consistency of the estimates across the four specifications of (DEM-regression) provides an empirical robustness check.

## 5 Results on the MEM

Our main results on the graph-based decomposition of drug identities into interpretable molecular descriptors are presented in this section. We consider the statistical model defined in (MEM-regression) and estimate the effects of the *P* = 57 molecular properties, together with all 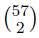 second-order interactions and 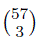 third-order interactions. This yields a regression model with up to

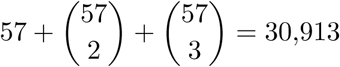

molecular effects. To position this high-dimensional regression model, a useful point of comparison is the statistical prediction of biochemical properties (i.e., toxicity, solubility, etc.) from graph-based molecular predictors. Our MEM formulation provides complementary evidence in a distinct regime: rather than predicting biochemical properties from structure alone, we ask whether drug response heterogeneity across pediatric AML samples can be explained by an explicitly constructed descriptor representation (and its higher-order interactions).

To address the high-dimensionality of our graph-based molecular predictors and their second-order and third-order interactions, we implement a sparse regression architecture based on penalized estimation and subsequent model refinement. We consider the ridge regularization problem:

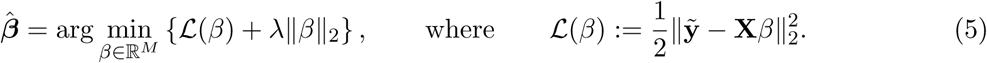

Here, **X** ∈ ℝ*^S^*^×*M*^ is the standardized design matrix collecting patient fixed effects, their interactions with log-concentration, and all main molecular effects and interactions up to order three, *S* is the number of point observations, *M >* 30,913 is the number of effects to be estimated, and **ỹ** ∈ ℝ*^S^* is the stacked vector of standardized responses. To approximate the minimizer ***β***^^^ in the high-dimensional regime, we use a Jacobi-type iteration. Specifically, define **A**: = **X**^⊤^**X** and **b**: = **X**^⊤^**ỹ**, let **D**: = diag(**A**) and **U**: = **A** − **D** denote the strictly off-diagonal residual operator. Starting from an initial ***β***^(0)^, we iterate

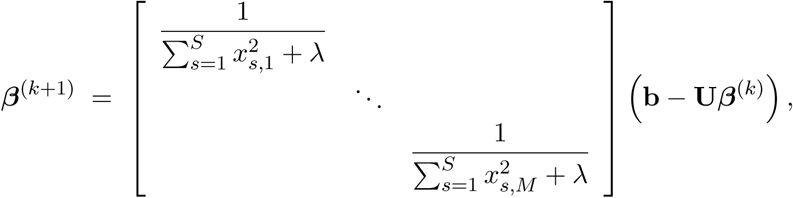

where *x_s,h_*denotes the (*s, h*) entry of **X** (the value of covariate *h* for observation *s*).

We apply this procedure to our data (Section 3). Figure 4 reports the log-scaled spectral decay of the top 100 ranked ridge regression coefficients. The pronounced elbow structure reveals a rapidly decaying spectrum, indicating strong compressibility of the molecular effect space and supporting the existence of a sparse underlying interaction structure.

**Figure 4:**
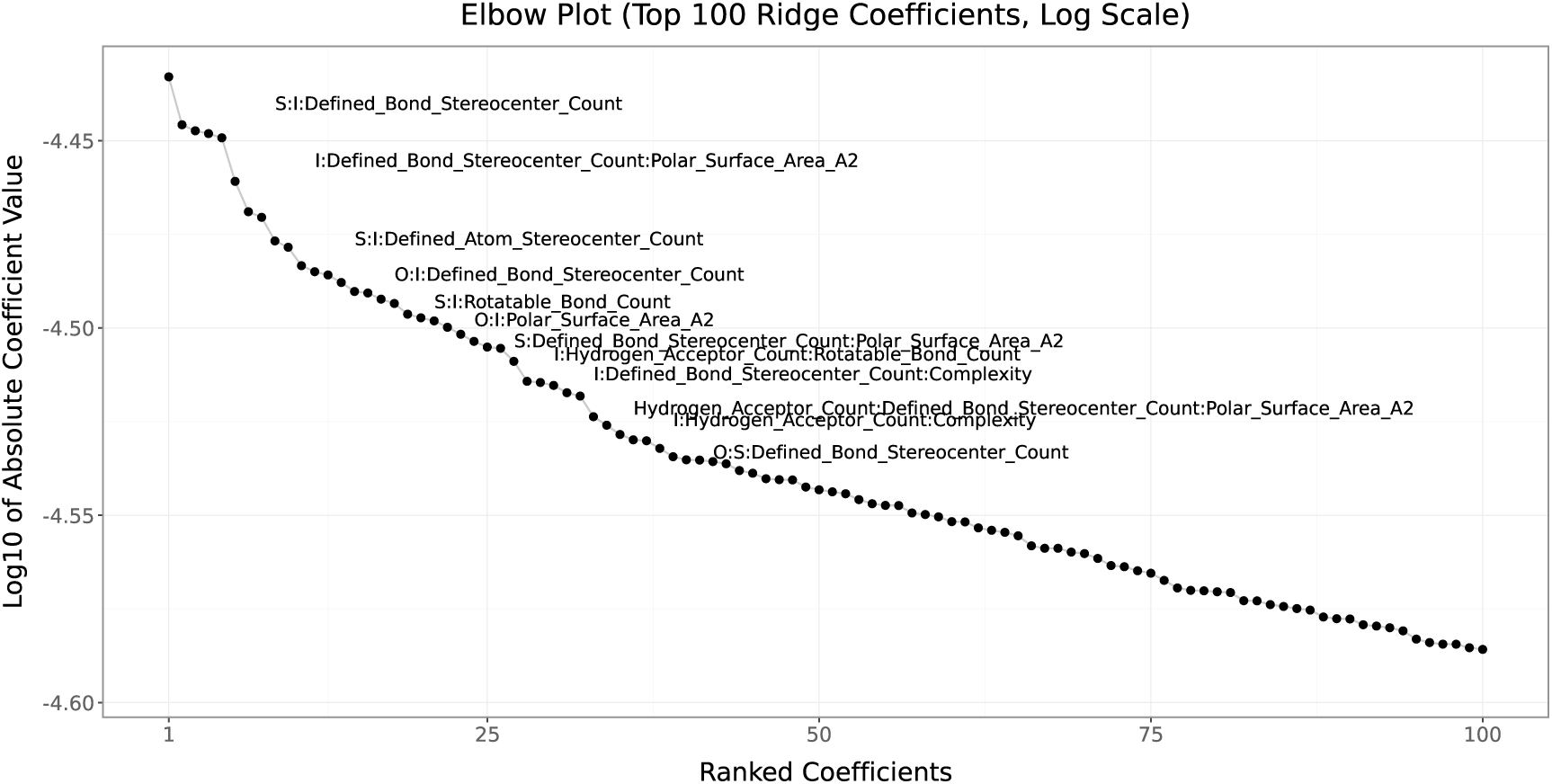
Log-scaled spectral decay of the top 100 ridge regression coefficients.

Since data are standardized, the result of this ridge allows ordering the *M* effects based on the absolute size of the ***β*** estimates, as illustrated in Figure 4. Guided by this ridge regression ranking, we proceed with a three-step iterative regression procedure designed to recover a statistically stable and interpretable MEM specification while excluding spurious high-order noise.

1. *(Forward selection)* Starting from an empty specification, terms are sequentially appended, following the descending order of the ridge regression ranking. A candidate term is retained if it satisfies all of the following criteria: (i) statistical significance at the 0.05 level, (ii) absence of multicollinearity (variance inflation factor (VIF) smaller than 10), and (iii) strict decrease in the Akaike Information Criterion (AIC). This step allows for the selection of terms *M̃* < *M*.
2. *(Pairwise exchange refinement).* In a second stage, we perform iterative pairwise exchanges between excluded and included terms. An exchange is accepted if (i) the entering term is statistically significant at *α* = 0.05, (ii) does not violate multicollinearity or rank conditions, and (iii) leads to a lower AIC than the incumbent specification. This step mitigates potential path-dependence introduced by greedy forward selection. This step does not alter the number of selected terms *M̃*.
3. *(Backward pruning).* Finally, we apply backward elimination to remove any remaining redundant terms, with the objective of obtaining the most parsimonious model that minimizes AIC. This step proceeds sequentially, and at each iteration it drops a non-significant term if the resulting regression has at least the same number of significant terms and at least the same AIC. This step enables the selection of terms *M̃* < *M̃*.

The combination of ridge regression ranking and the three-step refinement procedure allows us to isolate robust molecular effects while filtering out noise inherent to the combinatorial interaction space. This emphasis on linear-algebraic screening is aligned with the computational comparisons of Jiang et al. [2021a], who report that descriptor-based models (e.g., SVM, random forests, gradient boosting) typically train orders-of-magnitude faster than graph-based neural models, an important consideration when repeated fitting or resampling is required.

From this computational perspective, our methodology is in essence, related to classical step-wise regression procedures, which iteratively add and remove covariates based on a local criterion such as *p*-values or AIC. However, in our setting, a direct application of standard stepwise regression is impractical, as the candidate set comprises *M >* 30,913 highly dependent regressors (due to systematic overlap between molecular descriptors and their higher-order interactions). Our approach addresses these limitations by combining a global shrinkage-based screening stage with local refinement. Specifically, the ridge step allows creating a fully saturated model, which is not singular even in the presence of *M >* 30,913 highly dependent regressors. This produces a stable ordering of effects that act as a computationally tractable surrogate for exploring the full model space. Forward selection is then constrained to this ranked list and is coupled with explicit VIF and rank checks to prevent ill-conditioned updates, while the subsequent pairwise exchanges correct for path-dependence introduced by greedy inclusion. Finally, backward pruning removes redundant terms after joint refitting, yielding a parsimonious specification without sacrificing fit.

Applying our three-step procedure to our AML data set (Section 3), the final estimates of MEM retain only 45 molecular effects from the initial 30,913 possible candidates. Table 7 reports the estimated coefficients and corresponding *p*-values for the selected specification (patient fixed effects and patient-specific concentration effects are included but omitted from the table for readability). Despite the drastic reduction in dimension, the model achieves a strong explanatory performance, with marginal and conditional *R*^2^ values exceeding 0.61, which is close to the explanatory power of the corresponding DEM fits (Section 4).

**Table 7:**
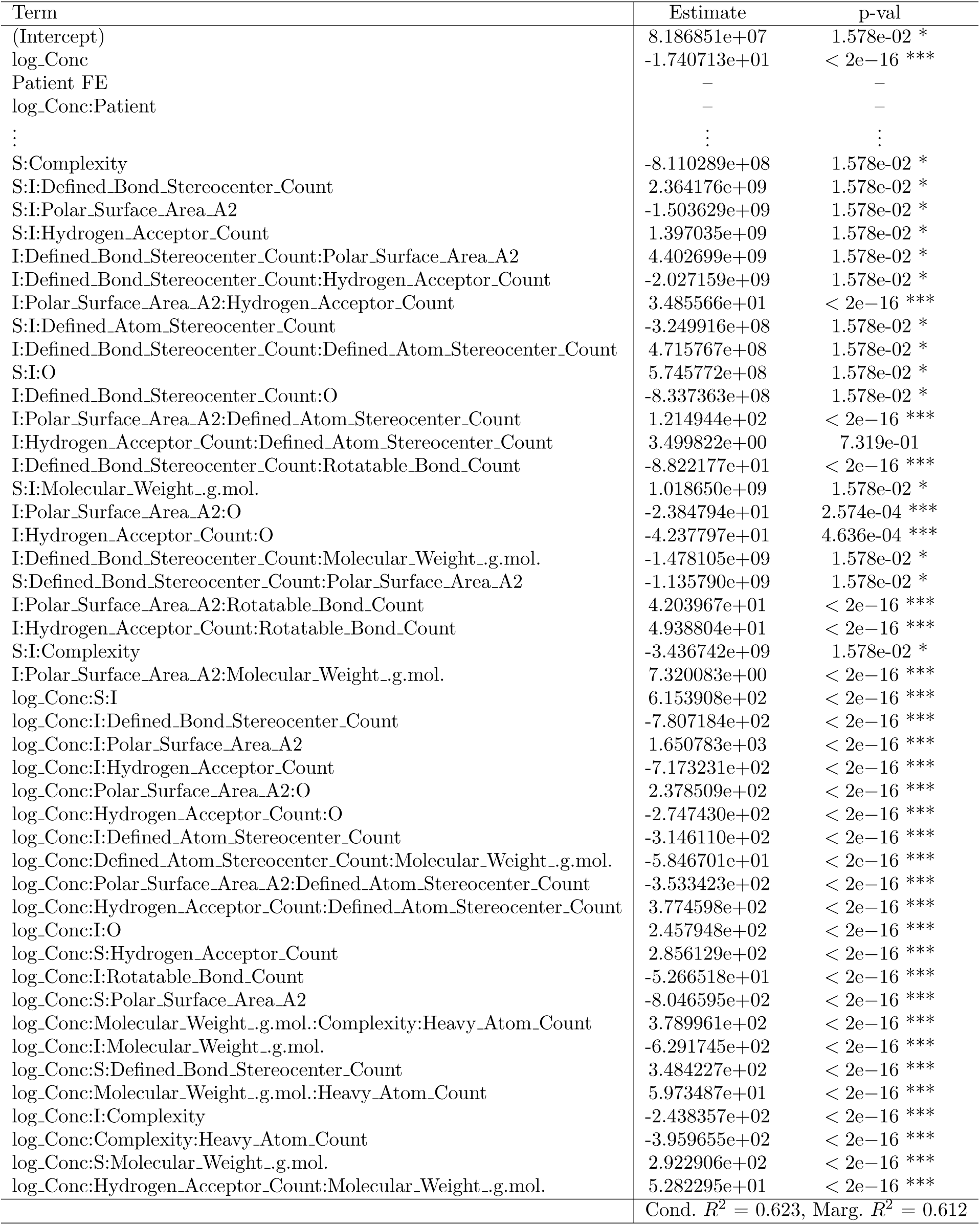
Zero-truncated viability score: model term estimates and p-values.

An important empirical finding is the uneven participation of molecular properties across retained interaction terms. Several descriptors appear recurrently in second- and third-order interactions, while many others are entirely excluded by the selection process. The coefficient on log Conc is strongly negative, confirming that dose escalation is the primary driver of decreased viability, consistent with the DEM dose–response patterns (Section 4 and Figure 3). Table 7 then shows that, once patient fixed effects and patient-specific concentration slopes are absorbed, the remaining explainable variation is concentrated in a small “physicochemical hub” of descriptors that repeatedly enters the retained interactions. In particular, Polar Surface Area A2, Hydrogen Acceptor Count, the presence of oxygen (O), stereochemical complexity (Defined_Atom_Stereocenter_Count and Defined_Bond_Stereocenter_Count), conformational flexibility (Rotatable_Bond_Count), and size proxies (Molecular_Weight, Heavy_Atom_Count, Complexity) appear systematically as second- and third-order terms and, crucially, as interactions with log_Conc. This pattern indicates that these features do not only shift baseline viability; rather, they also modulate the *slope* of the concentration–response relationship (i.e., how rapidly viability drops as dose increases) through the log_Conc:·interaction structure.

In particular, the prominence of polarity- and hydrogen-bond related quantities in our retained hub (e.g., Polar_Surface_Area_A2, Hydrogen_Acceptor_Count) echoes the post-hoc attribution patterns reported by Jiang et al. [2021a] for descriptor-based predictors, where SHAP analyses highlight polar and hydrophobicity-related descriptors as dominant drivers on representative end-points (e.g. solubility and BBB permeability). From a pharmacological standpoint, this convergence is consistent with a transport-and-engagement interpretation: polar surface area and hydrogen-bond acceptor capacity jointly influence aqueous solubility, membrane permeability, and the ability to form stabilizing contacts in binding pockets, while stereochemical descriptors capture the three-dimensional specificity of target engagement (enantiomeric/conformational selectivity). Likewise, the recurring involvement of rotatable bonds and global complexity/size measures suggests a flexibility–rigidity tradeoff. Crucially, however, the MEM yields an intrinsically parameterized and dose-conditional interpretation: through the signs and higher-order interaction structure (especially log_Conc:· terms), the model directly indicates whether these properties steepen or attenuate the concentration–response slope, thereby amplifying (synergistic sign) or dampening (antagonistic sign) dose-dependent cytotoxicity depending on the descriptor context.

Overall, this application of our hybrid three-step strategy has preserved the interpretability of stepwise selection of graph-based chemical properties, while remaining better suited to the combinatorial, interaction-rich MEM design than classical stepwise procedures applied directly to the full candidate set. Importantly, the fact that (MEM-regression) estimated based on our hybrid three-step strategy reaches *R*^2^ values close to the (DEM-regression) with only 45 retained molecular terms supports the central thesis of the paper: much of what the DEM attributes to “drug identity” can be reconstructed from a compact set of interpretable chemical attributes and their interactions. in concordance with Rudin [2019] and Jiang et al. [2021a], this fact reinforces the broader empirical message that carefully chosen descriptor representations can capture most of the signal that more complex structure-based representations aim to learn, here measured by the near-parity between MEM and DEM explanatory performance.

To help illustrate the results obtained, and in particular the uneven participation of molecular properties across retained interaction terms, Figure 5 provides a heatmap, representing the frequency with which each chemical property enters the retained interaction structure. Warmer colors indicate properties that play a central role in explaining drug viability outcomes.

**Figure 5:**
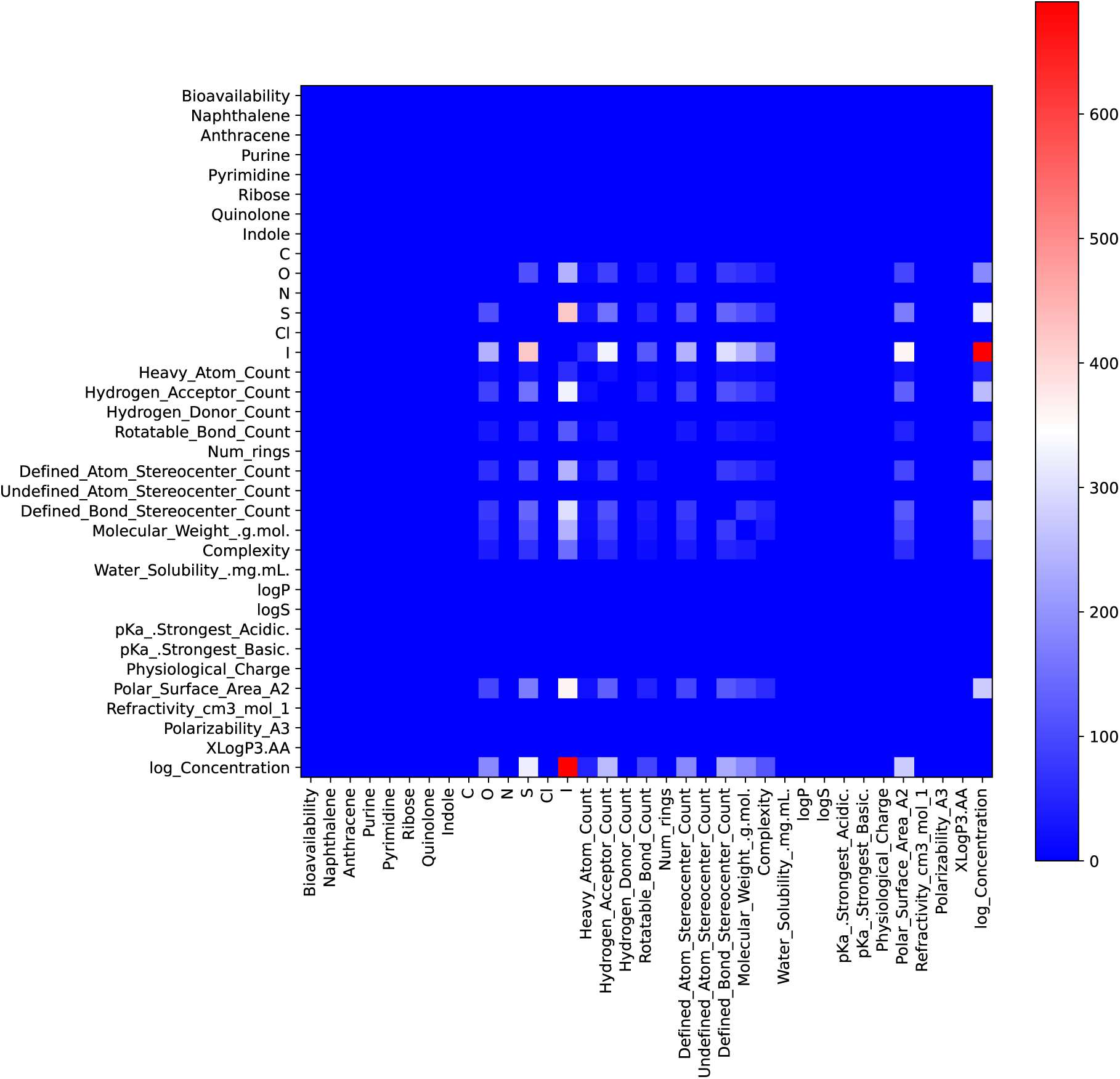
Heatmap illustrating the frequency of appearance of estimated interaction effects between chemical properties obtained from the zero-truncated MEM regression model

Beyond pairwise frequencies, the retained interaction structure naturally lends itself to a graph-based representation. Figure 6 depicts the resulting interaction network as a graph *G*(*V, E*), where the nodes *V* represent molecular properties and interaction terms. Molecular property nodes are shown as light blue circles when retained by (MEM-regression), and in gray otherwise, while third-order interaction terms are represented by square nodes. Edges connect chemical properties involved in second-order interactions, and third-order interaction nodes are linked to each of their constituent properties. The color of the interaction nodes is red or green according to the sign of the estimated coefficient, thereby encoding whether the corresponding interaction exerts an antagonistic or synergistic effect on drug viability.

**Figure 6:**
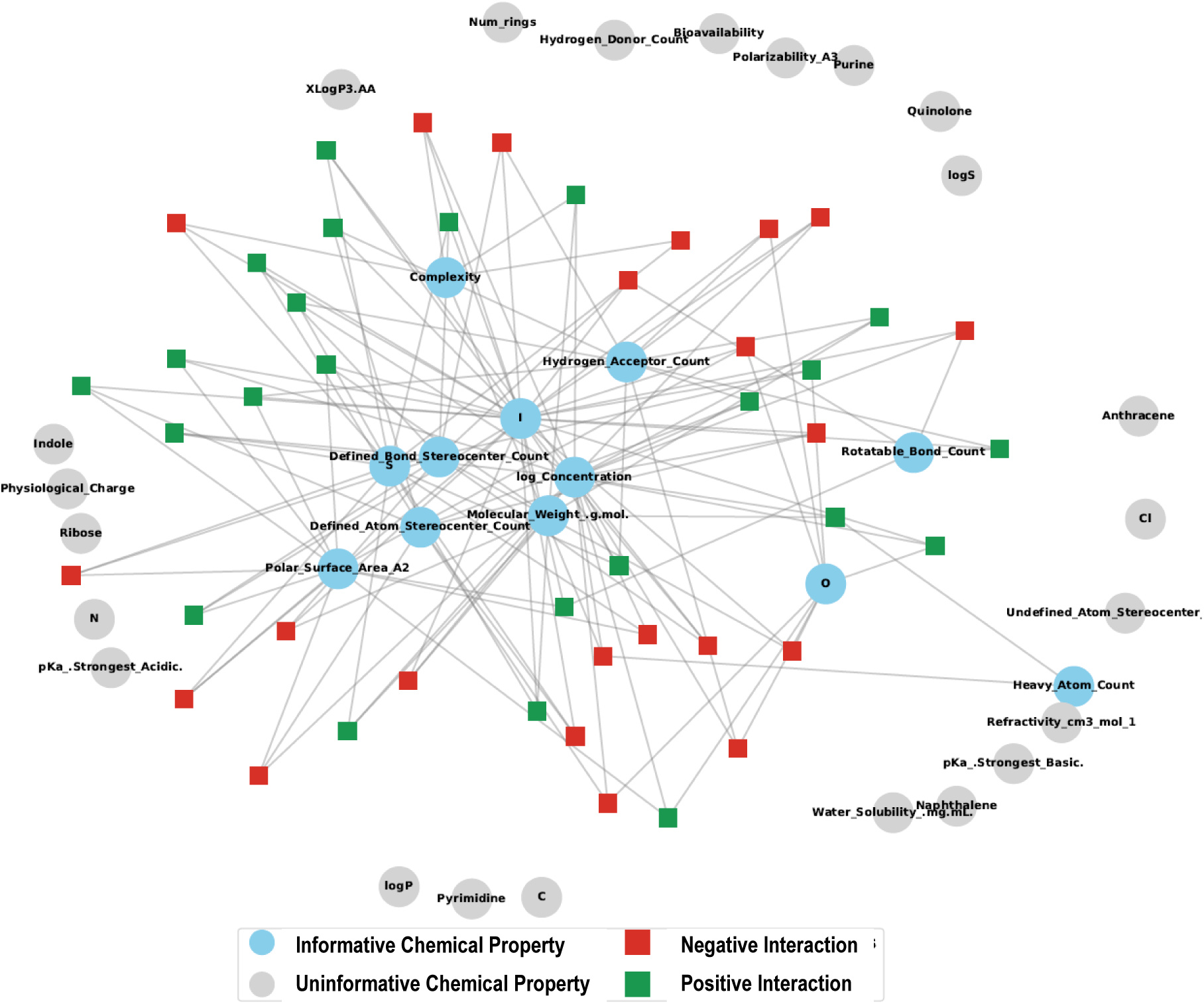
Visual representation of the graph network of estimated interactions

Taken together, these representations reveal a sparse yet highly structured interaction topology in which a small subset of molecular descriptors acts as hubs driving higher-order synergistic and antagonistic effects. Whereas Jiang et al. [2021a] demonstrate interpretability primarily through post-hoc explanation of black-box predictors (e.g., SHAP for gradient boosting), MEM produces a sparse, explicitly structured set of main and higher-order molecular effects that can be visualized as an interaction graph *G*(*V, E*) and used directly for downstream tasks such as principal compound or combination selection. Beyond interpretability, this graph-based structure provides a compact and operational representation of the molecular effect space, effectively summarizing the information contained in the estimated MEM coefficients. In the next subsection, we use *G*(*V, E*) to formulate a Molecular Effect Multi-Drug Selection Problem, where the estimated main and interaction effects from the (MEM-regression) directly guide the optimal selection and weighting of drug combinations.

### 5.1 Graph-based Multi-drug Selection Problem

This section shows how the estimated molecular interactions in the (MEM-regression) (see their representation as an interaction graph in Figure 6) provide a quantitative description of drug complementarity that can be exploited to guide the selection of candidate drug synergies. As noted by Al-Lazikani et al. [2012], drug combination discovery has often relied on ad hoc strategies, based primarily on empirical trial-and-error experimentation, an approach that becomes increasingly infeasible as the combinatorial complexity of multi-drug possibilities grows.

Motivated by this perspective, we introduce the Graph-based Multi-drug Selection Problem (GMSP), whose goal is to identify a combination of drugs from a set D of candidate drugs to minimize the predicted viability outcome. The objective explicitly accounts for individual molecular properties, pairwise interactions, and third-order interaction effects, as estimated by the three-step refinement procedure applied to the (MEM-regression). In what follows, we formally define a constrained ternary optimization model that captures this problem and show the numerical results obtained by applying the GMSP to the set of drugs reported in Table 4, using the biochemical covariates and interaction structure inferred from the MEM. To do so, we define G_1_ ⊆ P (with |G_1_| = *G*_1_), G_2_ ⊆ P × P (with |G_2_| = *G*_2_), and G_3_ ⊆ P × P × P (with |G_3_| = *G*_3_) as selected subsets of molecular properties and their second and third-order interactions, respectively. These sets are obtained from the empirical estimates based on the three-step refinement procedure summarized in Table 7. The GMSP relies on the following decision variables:

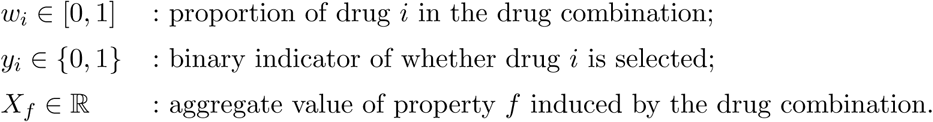

The model parameters (known quantities) are: *p_if_*, the value of the chemical property *f* associated with the drug *i*; *β*_0_, the estimated intercept; *β_f_*, the estimated coefficient of the property *f*; *β_fk_*, the estimated interaction coefficient between *f* and *k*; *β_fkl_*, the estimated interaction coefficient between *f*, *k*, and *l*; *N*, the targeted maximum number of drugs to be included in the combination; and *W*_min_, the minimum combination proportion assigned to any selected drug. The GMSP is defined as follows:

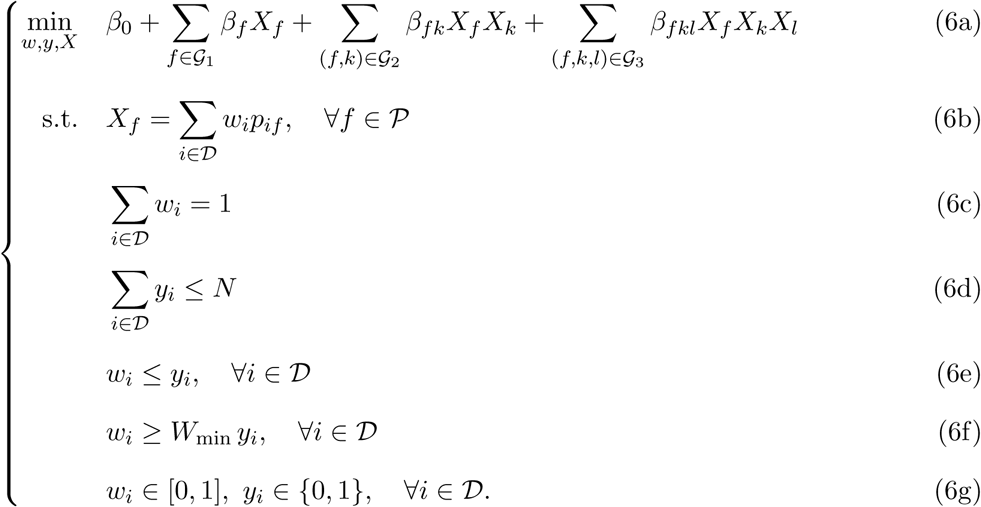

The objective function (6a) minimizes the predicted leukemic cell viability of the drug combination as estimated by the MEM. It consists of the intercept term *β*_0_, the contribution of selected main molecular effects *β_f_ X_f_* for all *f* ∈ G_1_, as well as second- and third-order interaction terms captured by *β_fk_X_f_ X_k_* and *β_fkl_X_f_ X_k_X_l_* for (*f, k*) ∈ G_2_ and (*f, k, l*) ∈ G_3_, respectively. This objective explicitly incorporates the high-order chemical complementarities identified in the (MEM-regression). Constraint (6b) defines combination-level aggregation of molecular properties: for each *f* ∈ P, the aggregate value *X_f_*is computed as the weighted sum of property values *p_if_* across candidate drugs, with weights given by the combination proportions *w_i_*. Constraint (6c) enforces that the combination proportions form a convex combination, ensuring that the selected proportions sum to one and thus define a valid drug synergy. Constraint (6d) controls combination cardinality by imposing that at most *N* drugs are selected, as indicated by the binary variables *y_i_*. Constraint (6e) links continuous proportions to the binary selection decisions by ensuring that a drug can receive a positive weight only if it is selected; in particular, if *y_i_* = 0, then *w_i_* = 0. Finally, constraint (6f) ensures that if a drug is included in the combination, it receives at least the minimum proportion *W*_min_.

Finally, we note that the objective function (6a) contains both quadratic and third-order interaction terms in the continuous variables *X_f_*, which renders the GMSP a nonconvex polynomial optimization problem. To enable tractable computation using standard mixed-integer optimization solvers, these nonlinear terms are reformulated via linearization. In particular, bilinear terms of the form *X_f_ X_k_* are replaced by auxiliary variables, say *Z_fk_*, together with a set of McCormick envelope constraints that characterize the convex hull of the bilinear relation when *X_f_*and *X_k_* are bounded. Given bounds 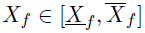 and 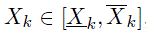, the equality *Z_fk_* = *X_f_ X_k_* can be enforced using four linear inequalities defining the McCormick relaxation.

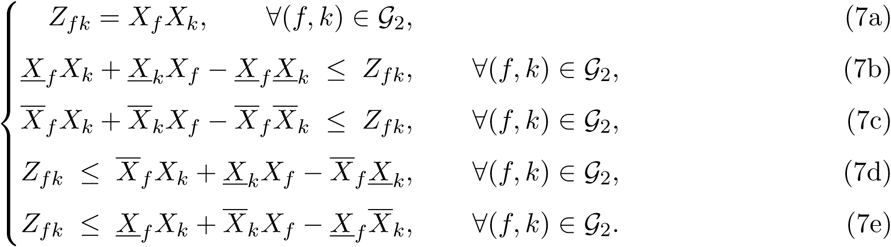

Third-order interaction terms of the form *X_f_ X_k_X_l_* are handled using a recursive linearization approach. Specifically, an auxiliary variable *Z_fk_* = *X_f_ X_k_* is first introduced and linearized via McCormick envelopes, after which a second auxiliary variable *T_fkl_* = *Z_fk_X_l_* is defined and linearized in the same manner. This sequential reformulation transforms the original polynomial objective into a linear function of the original and auxiliary variables, at the cost of additional constraints. As a result, the GMSP can be cast as a mixed-integer linear optimization problem with a well-defined feasible region, allowing the optimal drug combination to be computed efficiently using off-the-shelf solvers.

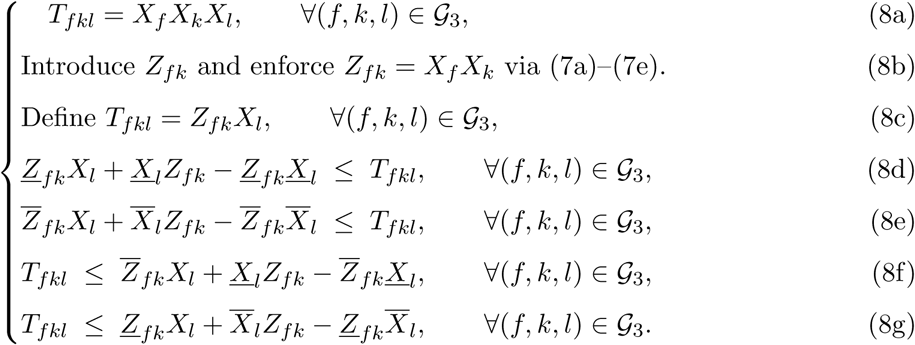

To solve the GMSP, we directly leverage the estimated 45 significant coefficients obtained from the (MEM-regression), including main effects, as well as second- and third-order interaction terms. The optimization model is instantiated using the set of 23 drugs previously evaluated in the (DEM-regression), ensuring full consistency between the predictive and prescriptive stages of the analysis. We solve the GMSP for varying combination cardinalities, ranging from *N* = 2 to *N* = 10, thereby exploring how optimal drug combinations evolve as the allowable size of the drug combination increases.

All optimization instances are solved using the quadratic optimization solver of Gurobi, implemented through its Python interface gurobipy (version 13.0). The resulting optimal drug selections, combination proportions, and objective values obtained for different combination cardinalities are summarized in Table 8. These results are directly guided by the estimated molecular effects and interaction structure inferred from the MEM, providing a principled and computationally tractable approach to multi-drug combination design.

**Table 8:** Summary of optimal GMSP solutions for different combination cardinalities.

| $N$ | Selected drugs | Combination proportions | Objective |
| --- | --- | --- | --- |
| 2 | MYLO, ABT199 | (0.90, 0.10) | -1.310e+13 |
| 3 | MYLO, ABT199, MIDO | (0.80, 0.10, 0.10) | -1.115e+13 |
| 4 | DASA, GILT, PALBO, FLAVO | (0.10, 0.70, 0.10, 0.10) | -6.114e+11 |
| 5 | MIDO, DASA, GILT, PALBO, FLAVO | (0.60, 0.10, 0.10, 0.10, 0.10) | -7.841e+11 |
| 6 | MYLO, DNR, ETO, IDR, ABT199, MIDO | (0.50, 0.10, 0.10, 0.10, 0.10, 0.10) | -6.102e+12 |
| 7 | MYLO, DNR, ETO, IDR, ABT199, NUTLIN, MIDO | (0.40, 0.10, 0.10, 0.10, 0.10, 0.10, 0.10) | -4.717e+12 |
| 8 | MYLO, DNR, ETO, IDR, ABT199, PONA, NUTLIN, MIDO | (0.30, 0.10, 0.10, 0.10, 0.10, 0.10, 0.10, 0.10) | -3.491e+12 |
| 9 | MYLO, DNR, ETO, IDR, ABT199, PONA, PRIMA, NUTLIN, MIDO | (0.20, 0.10, 0.10, 0.10, 0.10, 0.10, 0.10, 0.10, 0.10) | -2.446e+12 |
| 10 | MYLO, DNR, ETO, IDR, ABT199, PONA, PRIMA, NUTLIN, QUIZA, MIDO | (0.10, 0.10, 0.10, 0.10, 0.10, 0.10, 0.10, 0.10, 0.10, 0.10) | -1.567e+12 |

Inspection of Table 8 reveals a clear pattern of diminishing returns as the combination cardinality increases beyond two to three drugs. While the inclusion of a small number of complementary compounds yields substantial improvements in the predicted viability reduction, expanding the number of drugs further leads to progressively smaller gains. This behavior is consistent with the underlying chemical structure of the drug set: many candidate molecules share overlapping molecular properties and interaction profiles, allowing two or three drugs to effectively exploit chemical complementarity. Beyond this point, additional drugs primarily introduce redundancy, contributing limited new interaction effects and potentially adding noise and inefficiencies rather than enhancing the overall therapeutic impact.

## 6 Conclusion

This paper proposes an interpretable, graph-informed statistical framework for modeling *ex vivo* drug response in pediatric AML and for translating estimated molecular effects into actionable compound selection. Using viability measurements from 323 pediatric AML exposed *ex vivo* to up to 23 antileukemic drugs over replicated serial dilutions, we first quantify dose–response patterns with patient-aware regression models that capture both concentration effects and heterogeneous drug efficacy.

Our methodological contribution then consists in moving beyond drug identities as indivisible labels by decomposing compound effects into curated molecular properties and their higher-order interactions. Within the Molecular Effect Model (MEM), drug effects are expressed as linear combinations of interpretable chemical descriptors, allowing response variation to be attributed to specific molecular constituents rather than to opaque compound-level fixed effects. Although the full MEM expansion yields a combinatorial interaction space, our penalization-and-refinement strategy recovers a compact specification that retains only a small subset of robust molecular terms while achieving explanatory power close to the drug-identity baseline. Empirically, the retained interaction structure concentrates around a small physicochemical “hub” (notably polarity, hydrogen-bonding capacity, stereochemical complexity, flexibility, and size proxies), and its frequent interaction with concentration indicates that these properties primarily modulate the slope of the concentration–response relationship rather than merely shifting baseline viability. In line with the broader interpretability message emphasized in Jiang et al. [2021a] and Rudin [2019], these results illustrate that carefully structured descriptor representations can preserve predictive signal while enabling intrinsic, parameter-based interpretation.

Finally, we show how the estimated MEM interaction topology can be operationalized for down-stream decision-making. By representing retained main and interaction effects as a graph and embedding them into a Graph-based Multi-drug Selection Problem (GMSP), we obtain a prescriptive optimization layer that selects drug combinations by explicitly exploiting estimated chemical complementarities. The resulting solutions exhibit a clear pattern of diminishing returns beyond small combination sizes, suggesting that a limited number of complementary compounds can capture most of the attainable benefit within the considered candidate set, while additional drugs primarily introduce redundancy.

Several extensions follow naturally. On the modeling side, future work could incorporate richer patient-level biology (e.g., genomic or single-cell covariates) to connect molecular mechanisms of drug action to patient subtypes, and could explore alternative nonlinear concentration–response parameterizations while retaining intrinsic interpretability. On the validation side, the proposed decomposition-and-selection pipeline should be assessed on external cohorts and, ultimately, in prospective orthogonal evidence studies that evaluate whether the MEM-guided combinations yield improved *ex vivo* or clinical response. More broadly, the framework illustrates a path from interpretable molecular effect estimation to principled compound prioritization, offering a transparent alternative to purely black-box predictors for clinically oriented preclinical decision support.

## A Regression Diagnostics

Table A.1 indicate that the adjusted collinearity measures *GV IF* ^1*/*(2*Df*)^ are about 1.04–1.31 for all blocks (Drug, Patient, and their interactions with log-concentration) in both the zero-truncated and bounded viability models. These values are close to 1 and well below common concern thresholds (e.g., 2–5), indicating no practically relevant multicollinearity among the factor blocks or their interactions. The very large raw GVIF numbers reflect the no-intercept parameterization and the high dimensionality of the dummy sets.

**Table A. 1:** Generalized variance–inflation diagnostics for two specifications of the viability outcome in Table 4. Entries report *GV IF*, the associated degrees of freedom (df), and the scale-free quantity *GV IF* ^1^*^/^*^(2^*^Df^*^)^ that is comparable across multi-df factor blocks. Since *GV IF* and *GV IF* ^1^*^/^*^(2^*^Df^*^)^ depend only on the design matrix, not on the response variable *V_i,j_*, the same multicollinearity analysis is valid for both zero-truncated viability scores and bounded viability scores. Lower values indicate weaker collinearity; as a rule of thumb, *GV IF* ^1^*^/^*^(2^*^Df^*^)^ *<* 2 is generally acceptable.

| Term | GVIF | df | $GVIF^{1/(2Df)}$ |
| --- | --- | --- | --- |
| Drug | $7.900424 \times 10^6$ | 38 | 1.232419 |
| Patient | $1.4351121 \times 10^8$ | 289 | 1.033028 |
| Patient:log-Concentration ($\lambda_{i,\cdot}^{LC}$) | $8.329734784 \times 10^9$ | 290 | 1.040171 |
| Drug:log-Concentration ($\lambda_{\cdot,j}^{LC}$) | $1.98399961 \times 10^8$ | 38 | 1.285814 |

## Footnotes

1 https://www.conect-aml.fr

2 As demonstrated by us and others [Tyner et al., 2018, Bottomly et al., 2022b, de Haar-Holleman et al., 2004, Gonzales et al., 2022], *ex vivo* drug resistance allows minimal alteration of natural conditions, while robustly isolating the drug target and disease-relevant features from physiological confounding interactions.

